# Subset selection of markers for genome-enabled prediction of genetic values using radial basis function neural networks

**DOI:** 10.1101/490474

**Authors:** Isabela de Castro Sant’ Anna, Gabi Nunes Silva, Moysés Nascimento, Cosme Damião Cruz

## Abstract

This paper aimed to evaluate the efficiency of subset selection of markers for genome-enabled prediction of genetic values using radial basis function neural networks (RBFNN). For this purpose, an F1 population from hybridization of divergent parents with 500 individuals genotyped with 1,000 SNP-type markers was simulated. Phenotypic traits were determined by adopting three different gene action models – additive, additive-dominant, and epistasic, complying with two dominance situations: partial and complete with quantitative traits admitting heritability (h^2^) equal to 30 and 60%, each one controlled by 50 loci, considering two alleles per locus, totaling 12 different scenarios. To evaluate the predictive ability of RR_BLUP and the neural networks, a cross-validation procedure with five replicates were trained using 80% of the individuals of the population. Two methods were used: dimensionality reduction and stepwise regression. The square of the correlation between the predicted genomic estimated breeding value (GEBV) and the phenotype value was used to measure predictive reliability. For h^2^ = 0.3 in the additive scenario, the R^2^ values were 59% for neural network (RBFNN) and 57% for RR-BLUP, and in the epistatic scenario, R^2^ values were 50% and 41%, respectively. Additionally, when analyzing the mean-squared error root, the difference in performance between the techniques is even greater. For the additive scenario, the estimates were 91 for RR-BLUP and 5 for neural networks and, in the most critical scenario, they were 427 for RR-BLUP and 20 for neural network. The results showed that the use of neural networks and variable selection techniques allows capturing epistasis interactions, leading to an improvement in the accuracy of prediction of the genetic value and, mainly, to a large reduction of the mean square error, which indicates greater genomic value.

## Introduction

One of the major challenges of genetic breeding today is understanding the genetic variation of quantitative traits, QTL (Quantitative trait loci), which are conditioned by a large number of genes with small effects [1] whose interaction often results in non-linearity in relations between phenotypes and genotypes [2,3].

With the advent of Genomic Selection (GS) [4], it became possible to estimate the genomic value of individuals (GEBV) without the need of phenotyping, which led to an increase in genetics gain by reducing time and money. Therefore, for many traits of agronomic importance, genetic values are determined by multiple genes of small effects and their phenotypic expression is strongly affected by genetic interaction between their additive, dominant and epistatic effects. However, most applications of GS include only the additive portion of the genetic value, so a more realistic representation of the genetic architecture of quantitative traits should include dominance and epistatic interactions, since these effects are crucial factors to increase the accuracy of prediction [5].

The inclusion of these interactions is computationally challenging and leads to the superparametrization of the models that are already in high dimensionality because of the large number of markers in the genome and the smallest number of individuals [2]. Besides that, before fitting the model it is necessary to define the model effects to be estimated. In this context, Artificial Neural Networks (ANNs) have a great potential because they can capture non-linear relationships between markers from the data themselves (without a previous model definition), which most of the models commonly used in the GS cannot do [2,6,7].

Radial Basis Function Neural Networks (RBFNN) are a particular class of Neural Networks (NN) that have properties that make it attractive to GS applications. According to Gianola et al. [8], RBFNNs have the ability to learn from the data used in their training, have universal approximation properties [9], give a unique solution and are faster than standard ANNs [10].

However, the inclusion of all markers in the prediction RBFNN model increases the chances of a high correlation between the markers [11]. The number of markers represents a huge challenge that leads to less precision and a great computational demand for NN training. This happens because NNs use good part of their resources to represent irrelevant portions of the search space and compromising the learning process because there are thousands of markers available in the genome [12]. Thus, a more realistic model should include only SNPs related to the traits of interest [6].

For this reason, a subset of SNPs can be used for training, since, by reducing the search space, RBFNNs improve the learning process and increase the predictive power of the model, as realized by [2]. These authors used two types of RBFNN models: one considering a common weight parameter for each SNP, and the other in which each SNP has specific parameters of importance. However, due to the importance of NNs for the improvement of prediction of quantitative traits, there is still a need to test different dimensionality reduction methods and prediction models for polygenic traits.

In view of the above, this paper aimed to evaluate the efficiency of genome-enabled prediction by radial basis function neural networks (RBFNN) in the prediction of genetic values by considering a subset of markers in simulated data set with different gene actions (as dominance and epistasis) and degrees of heritability. The results were compared with those obtained by one of the standard GS model: RR-BLUP.

## Material and methods

### Origin of populations

In order to assess the reliability of GS predictions, data were simulated by considering a diploid species with 2n = 2x = 20 chromosomes as the reference, and the total length of the genome was stipulated in 1.000 cM. Genomes were generated with a saturation level of 101 molecular markers spaced by 1cM per linkage group, totaling 1010 markers. Divergent parental line genomes were simulated, as well as genomes from the base population (F_1_). Since the base population was derived from two contrasting homozygous parents, the effective size of the base population is the size of F_1_ itself.

### Simulation of quantitative traits

Quantitative traits were simulated in three scenarios by considering three degrees of dominance (d/a = 0, 0.5 and 1) and two broad sense heritability (h^2^ = 0.30 and 0.60), considering two gene actions: additive and epistatic, thus totaling six scenarios. Each trait was controlled by 50 randomly chosen loci, with 2 alleles per locus.

The phenotypic values of the i^th^ individuals were obtained according to the additive model as follows:

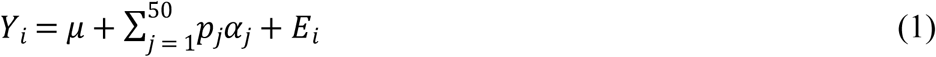

where *α*_*j*_ is the effect of the favorable allele in locus *j*, considered equal to 1, 0 or −1 for the genotypic classes AA, Aa and aa, respectively, and *p*_*j*_ is the contribution of locus *j* to the manifestation of the trait under consideration. In this study, the contribution of each locus was established as being equivalent to the probability of the set generated by the binomial distribution X∼ b (a+b)^s^, where a=b=0.5 and s = (50). The value of d_i_ was defined according to the average degree of dominance expressed in each trait. *E*_*i*_ is the environmental effect, generated according to a normal distribution with means equal to zero and variance given by the equation bellow:

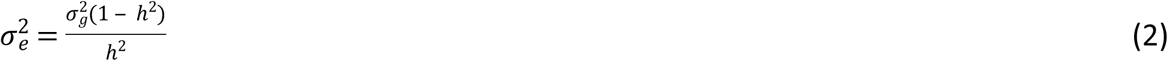

Where 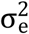 is the variance given by the environmental values, 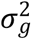 is the variance of the genetic values, and *h*^2^ is the heritability defined for the trait. The genetic variance is defined from the information of the genetic control and the importance of each locus in the polygenic model.

For the epistatic model, the phenotypic values of the i^th^ individuals were obtained according to the following equation:

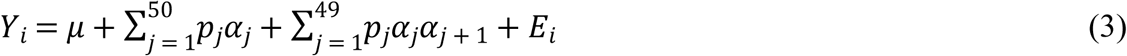

In the above equation, the first summation of the expression refers to the contribution of the individual locus through its additive and dominant effects and the second summation represents the multiplicative effects corresponding to the epistatic interactions between pairs of loci.*α*_*j*_ is the multiplicative effect of the favorable allele in locus *j*, and *j+1* and *p*_*j*_ is the contribution of locus *j* to the manifestation of the trait under consideration.

## RR-BLUP

The RR-BLUP model was used to obtain the genomic estimated breeding values (GEBV) [4]:

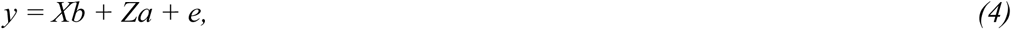

where *y* is the vector of phenotypic observations, *b* is the vector of fixed effects, *a* is the vector of random marker effects, and *e* refers to the vector of random errors, 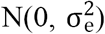; *X* and *Z* are matrices of incidence for *a* and *b*. The effects of the individuals (GEBVs) were estimated by the equation below:

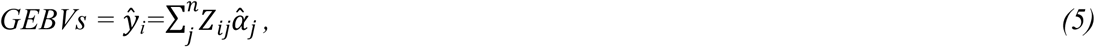

where *n* is the number of markers arranged in the genome, Z_ij_ is the line of the matrix of incidence that allocates the genotype of the j^th^ marker for each individual (i), 1, 0, −1 for genotypes A_1_A_1_, A_1_A_2_, A_2_A_2_, respectively, for biallelic and codominant markers, and 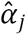 is the effect of the j^th^ marker estimated by RR-BLUP. In this model, the incidence matrix associated with the effects of dominance was not included. However, it should be remembered that the population has probably allele frequency being *p* different from *q* and therefore the additive effects estimated through matrix *Z* capture dominance effects.

### Radial Basis Function Neural Network (RBFNN)

A RBFNN is an artificial neural network that uses radial basis functions as activation functions. The RBFNN in the present study is a three layered feed-forward neural network, where the first layer is linear and only distributes the input signal, while the next layer is nonlinear and uses Gaussian functions (Fig 1).

**Fig. 1.**
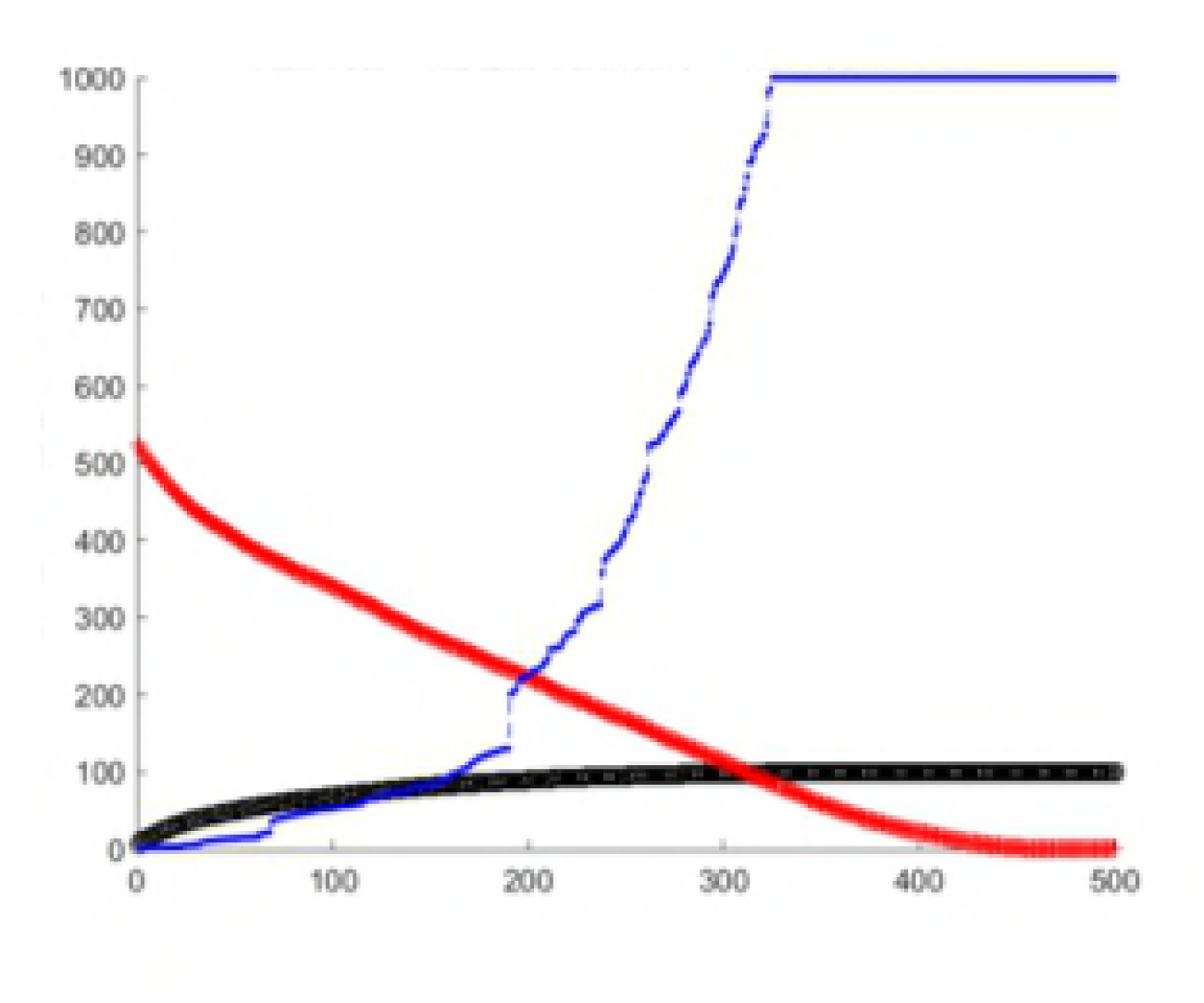
Structure of a radial basis function neural network (RBFNN). In the hidden layer, each input vector (**x_i1_**,…,**x_ip_**)is summarized by the Euclidean distance between the input vectors xi and the centers c_m_,(m = 1, …, M) neurons, i.e., h_m_||x_i_ – c_m_||, where hm is a bandwidth parameter. Then distances are transformed by the Gaussian kernel **exp**(-(h_m_||xi – c_m_||)2) for obtaining the response, 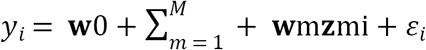 (extracted from Gonzalez-Camacho et al., 2012).

The training of RBFNN optimization includes: the weights between the hidden layer and the output layer, the activation function, the center of activation functions, the distribution of center of activation functions, and the number of hidden neurons [13]. During the training process, only the weights between the hidden layer and the output layer are modified. The vector of weights ω ={*W*_1_, …, *W*_*s*_} of the linear output layer is obtained using the ordinary least-squares fit that minimizes the mean squared differences between _yi_ (from RBFNN) and the observed _yi_ observed in the training set, provided that the Gaussian RBFs for centers c_k_ and h_k_ of the hidden layer are defined.

The radial basis function selected is usually a Gaussian kernel selected using K-means clustering algorithm. The centers are selected using the orthogonalization least-squares learning algorithm, as described by [14] and implemented in [15]. The centers are added iteratively such that each new selected center is orthogonal to the others. The selected centers maximize the decrease in the mean squared error of the RBFNN, and the algorithm stops when the number of centers (neurons) added to the RBFNN attains a desired precision (goal error) or when the number of centers is equal to the number of input vectors, that is, when S=n.

To select the best RBFNN, a grid for training the net was generated, containing different spread values and different precision values (goal error). The spread value ranging from 5 to 100 and an initial value of 0.01 for the goal error was considered. The spread parameter allows adjusting the form of the Gaussian RBFNN such that it is sufficiently large to respond to overlapping regions of the input space but not so big that it could induce the Gaussian RBFNN to have a similar response [16].

### SNP subset selection

To determine the number of markers, stepwise regression was used in the scenario with epistatic effects, dominance and low heritability. In this procedure, the maximum number of markers was determined in conjunction with measures representative of the data as the mean square error root of the model (MSER), determination coefficient (R^2^) obtained by inclusion of the selected markers, and the condition number (CN) of the correlation matrix. As for the first two criteria, the MSER chosen was the one that presented the lowest possible value tied to the best possible values for R^2^ (the higher the better). The third criterion was used to avoid multicollinearity problems. The condition number of the correlation matrix between the explanatory variables verifies the degree of multicollinearity in the correlation matrix X’X [17]. When the CN resulting from this division was lower than or equal to 100, it was considered that there was weak multicollinearity between the explanatory variables; for 100 < CN <1000 moderate to severe multicollinearity, and for CN≥1000 severe multicollinearity was considered. So, based on a graphical analysis, the number was determined by the graphical point with the best R^2^, the lowest REQM when 100 <CN.

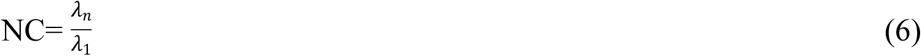

where *λ*_*n*_ is the eigenvalue of largest absolute value and *λ*_*1*_ of the smallest.

### Computational applications for data analysis

The models were compared using the reliability (R^2^) defined as the squared correlation between the predicted GEBVs of the individuals with no phenotypic traits and the root mean squared error (RMSE) using predicted and realized values. A five-fold cross-validation scheme was used to determine the reliability of genomic prediction of a selected subset of SNPs in the population. The individuals (500) were randomly split into five equal-size groups and each group with about 100 individuals (20% of the population) was in turn assigned with phenotypic values and used as the validation set. The reliability of genomic prediction was calculated as the squared correlation between the predicted GEBVs of the individuals with no phenotypic traits. The reliability reported in the study was the average of the reliability of genomic prediction from 5-fold groups. For comparison purposes, the reliability of genomic prediction from all the SNPs (1000) was also calculated, in addition to 100 SNPs selected to be even.. The simulations were implemented with software GENES [18] and the statistical analyses were performed with software R, with the RR-BLUP package [19].

## Results

Dimensionality reduction was performed using a graphical procedure that considers the of the model, the determination coefficient (R^2^) obtained by including the selected markers and the condition number (CN) of the correlation matrix. The number of markers was determined by the graphical point which presented the larger (R^2^ and the lowest MSER when 100 <CN (Fig 2). After defining the optimal number of markers, stepwise regression was used to select, among all markers, those used in the fit.

**Figure 2:**
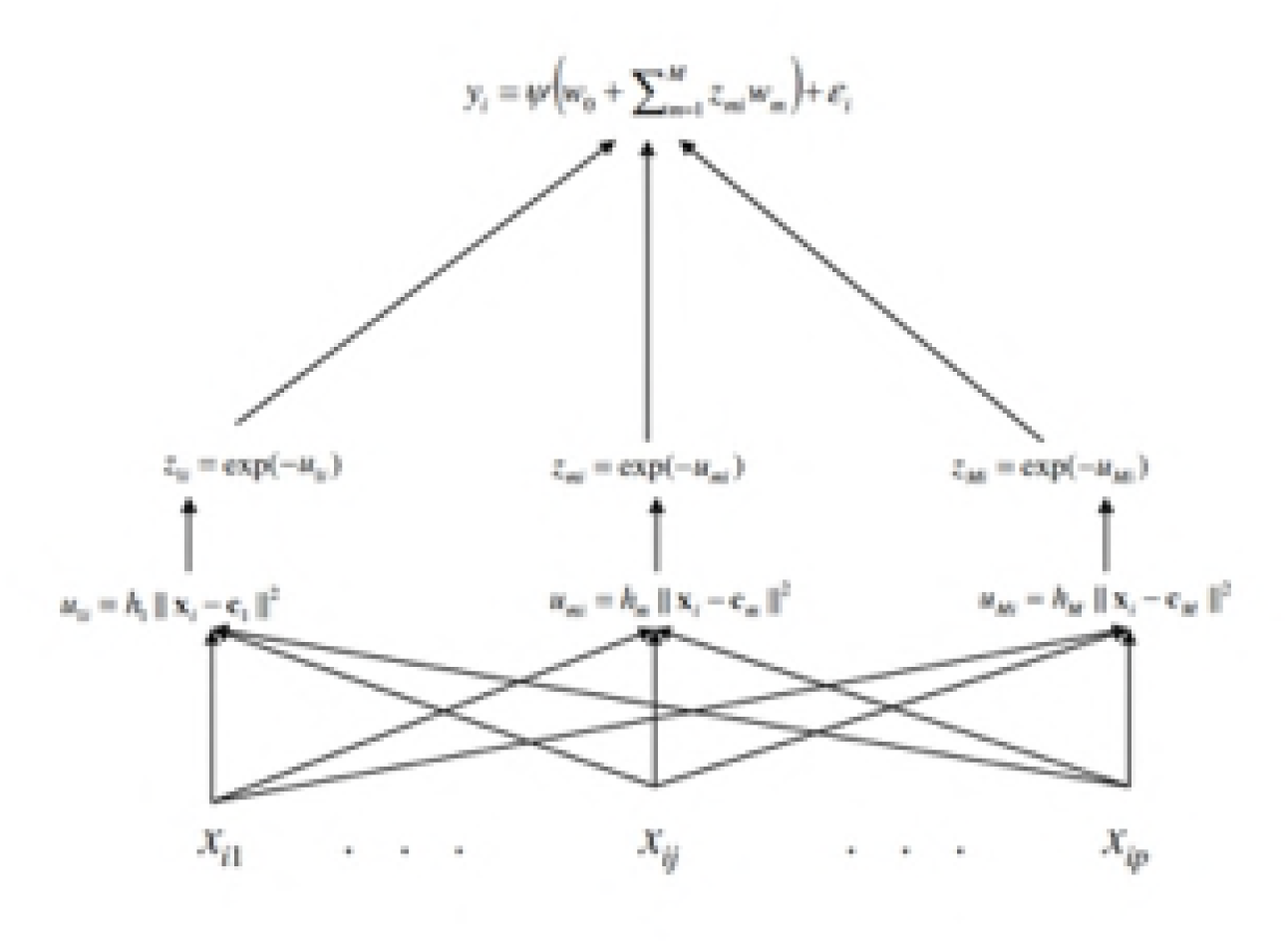
A graphical representation of the values of determination coefficient (R^2^) in black, Mean squared error root (MSER) in red and the condition number(CN) in blue obtained by the stepwise regression method by including 1 to 400 molecular markers (from the total of 1000) in the stepwise regression model.

Twelve different scenarios considering different levels of heritability, dominance and epistatic effects were evaluated (Table 2 and 3). Five cross-validation folds were used to access the reliability (R^2^) of fit models (RBFNN and RR-BLUP), considering or not dimensionality reduction.

**Table 1.**
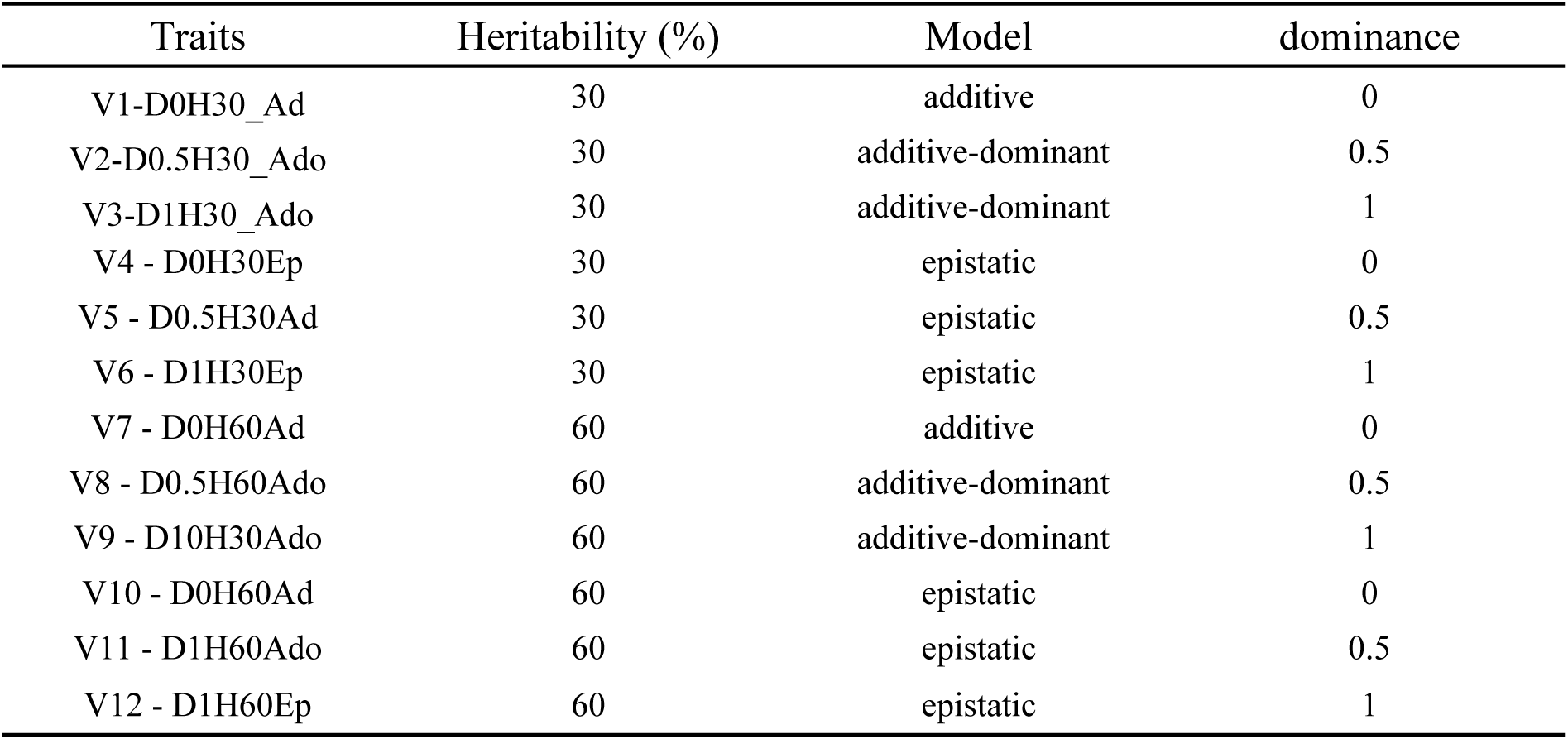
Scenarios composed by combination of traits, action genenic model. heritabilty and dominance degree.

| Traits | Heritability (%) | Model | dominance |
| --- | --- | --- | --- |
| V1-D0H30_Ad | 30 | additive | 0 |
| V2-D0.5H30_Ado | 30 | additive-dominant | 0.5 |
| V3-D1H30_Ado | 30 | additive-dominant | 1 |
| V4 - D0H30Ep | 30 | epistatic | 0 |
| V5 - D0.5H30Ad | 30 | epistatic | 0.5 |
| V6 - D1H30Ep | 30 | epistatic | 1 |
| V7 - D0H60Ad | 60 | additive | 0 |
| V8 - D0.5H60Ado | 60 | additive-dominant | 0.5 |
| V9 - D10H30Ado | 60 | additive-dominant | 1 |
| V10 - D0H60Ad | 60 | epistatic | 0 |
| V11 - D1H60Ado | 60 | epistatic | 0.5 |
| V12 - D1H60Ep | 60 | epistatic | 1 |

**Table 2.**
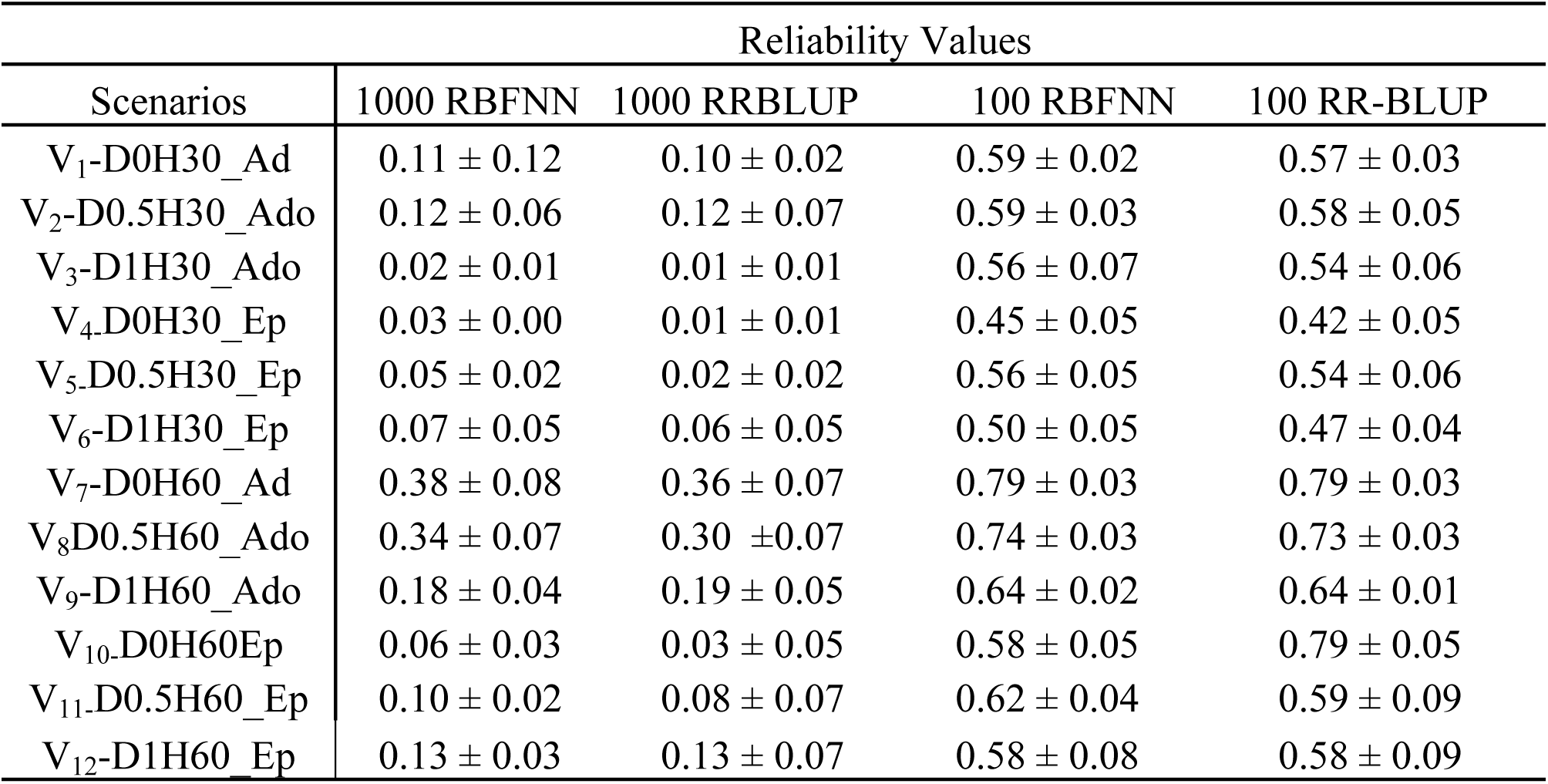
Reliability values of selection obtained from RBFNN (Radial Basic Neural Network) and RR-BLUP through all markers (1000) or selected markers (100) by Stepwise Regression(SWR) in a set of validation data involving cross-validation procedures.

| Scenarios | Reliability Values |  |  |  |
| --- | --- | --- | --- | --- |
|  | 1000 RBFNN | 1000 RRBLUP | 100 RBFNN | 100 RR-BLUP |
| V<sub>1</sub>-D0H30_Ad | 0.11 ± 0.12 | 0.10 ± 0.02 | 0.59 ± 0.02 | 0.57 ± 0.03 |
| V<sub>2</sub>-D0.5H30_Ado | 0.12 ± 0.06 | 0.12 ± 0.07 | 0.59 ± 0.03 | 0.58 ± 0.05 |
| V<sub>3</sub>-D1H30_Ado | 0.02 ± 0.01 | 0.01 ± 0.01 | 0.56 ± 0.07 | 0.54 ± 0.06 |
| V<sub>4</sub>-D0H30_Ep | 0.03 ± 0.00 | 0.01 ± 0.01 | 0.45 ± 0.05 | 0.42 ± 0.05 |
| V<sub>5</sub>-D0.5H30_Ep | 0.05 ± 0.02 | 0.02 ± 0.02 | 0.56 ± 0.05 | 0.54 ± 0.06 |
| V<sub>6</sub>-D1H30_Ep | 0.07 ± 0.05 | 0.06 ± 0.05 | 0.50 ± 0.05 | 0.47 ± 0.04 |
| V<sub>7</sub>-D0H60_Ad | 0.38 ± 0.08 | 0.36 ± 0.07 | 0.79 ± 0.03 | 0.79 ± 0.03 |
| V<sub>8</sub>-D0.5H60_Ado | 0.34 ± 0.07 | 0.30 ± 0.07 | 0.74 ± 0.03 | 0.73 ± 0.03 |
| V<sub>9</sub>-D1H60_Ado | 0.18 ± 0.04 | 0.19 ± 0.05 | 0.64 ± 0.02 | 0.64 ± 0.01 |
| V<sub>10</sub>-D0H60Ep | 0.06 ± 0.03 | 0.03 ± 0.05 | 0.58 ± 0.05 | 0.79 ± 0.05 |
| V<sub>11</sub>-D0.5H60_Ep | 0.10 ± 0.02 | 0.08 ± 0.07 | 0.62 ± 0.04 | 0.59 ± 0.09 |
| V<sub>12</sub>-D1H60_Ep | 0.13 ± 0.03 | 0.13 ± 0.07 | 0.58 ± 0.08 | 0.58 ± 0.09 |

**Table 3.** Mean squared error root obtained from RBFNN and RR-BLUP through all markers (1000) or selected markers (100) by Stepwise Regression(SR)in a set of validation data involving cross-validation procedures.

| MSER –mean squared error root |  |  |  |  |
| --- | --- | --- | --- | --- |
| Scenarios | 1000 RBF | 1000 RR-BLUP | 100 RBF | 100-RRBLUP |
| V<sub>1</sub>-D0H30_Ad | 5.9 ± 0.1 | 33.7 ± 2 | 4.9 ± 0.1 | 23.4 ± 0.0 |
| V<sub>2</sub>-D0.5H30_Ado | 6.2 ± 0.1 | 36.0 ± 1.3 | 5.1 ± 0.1 | 24.9 ± 3.7 |
| V<sub>3</sub>-D1H30_Ado | 16.8 ± 1.2 | 58.2 ± 5.7 | 13.5 ± 0.1 | 34.9 ± 17.7 |
| V<sub>4</sub>-D0H30_Ep | 16.9 ± 1.2 | 267.9 ± 10.5 | 14.2 ± 0.1 | 205.9 ± 48.1 |
| V<sub>5</sub>-D0.5H30_Ep | 18.5 ± 0.6 | 338.1 ± 17.7 | 15.5 ± 0.3 | 232.1 ± 18.9 |
| V<sub>6</sub>-D1H30_Ep | 23.8 ± 1.7 | 534.6 ± 24 | 20.8 ± 0.3 | 395.9 ± 40.6 |
| V<sub>7</sub>-D0H60_Ad | 4.5 ± 0.1 | 20.7 ± 1 | 3.4 ± 0.4 | 11.99 ± 0.9 |
| V<sub>8</sub>-D0.5H60_Ado | 4.7 ± 0.2 | 22.4 ± 2 | 3.7 ± 0.1 | 107.4 ± 1.1 |
| V<sub>9</sub>-D1H60_Ado | 5.4 ± 0.2 | 28.23 ± 2 | 4.3 ± 0.1 | 145.9 ± 5.2 |
| V<sub>10</sub>-D0H60Ep | 13.5 ± 0.3 | 181.5 ± 12 | 11.7 ± 0.2 | 320.1 ± 36.4 |
| V<sub>11</sub>-D0.5H60_Ep | 15.5 ± 0.5 | 236.8 ± 19 | 12.4 ± 0.3 | 280.9 ± 28.9 |
| V<sub>12</sub>-D1H60_Ep | 18.8 ± 0.8 | 355.5 ± 26 | 15.7 ± 0.6 | 473.8 ± 22.0 |

Overall, dimensionality reduction improved the reliability values for all scenarios, specifically, with h^2^ =30 the reliability value from 0.03 to 0.59 using RBFNN and from 0.10 to 0.57 with RR-BLUP in the scenario with additive effects. In the additive dominant scenario, the reliability values changed from 0.12 to 0.59 using RBFNN and from 0.12 to 0.58 with RR-BLUP, and in the epistasis scenarios the reliability values changed from 0.07 to 0.50 using RBFNN and from 0.06 to 0.47 with RR-BLUP (Table 2).

In the scenarios with h^2^=60, the reliability value improved from 0.38 to 0.79 using RBFNN and from 0.36 to 0.79 with RR-BLUP in the scenario with additive effects. In the scenario with additive dominance, the values changed from 0.34 to 0.79 using RBFNN and from 0.30 to 0.73 with RR-BLUP, and in the epistatic scenarios the average of reliability values changed from 0.10 to 0.60 using RBFNN and from 0.08 to 0.58 with RR-BLUP (Table 2).

Table 3 shows the range of values for the accuracy of prediction (MSER = mean squared err root). For all scenarios with and without dimensionality reduction, RBFNN outperformed RRBLUP. Besides that, dimensionality reduction also improved the accuracy of RBFNN and RRBLUP. The MSER value ranged from 3.5 to 23.8 for RBFNN and from 71.4 to 575.4 for RR-BLUP. Specifically, with h^2^ =30 the MSER value ranged from 5.9 to 4.9 using RBFNN and from 33.7 to 23.4 with RR-BLUP in the scenario with additive effects. In the additive-dominance scenario, the average of MSER values changed from 11.5 to 9.3 using RBFNN and from 47.1 to 29.4 with RR-BLUP, and in the epistasis scenarios the average of MSER values changed from 19.73 to 13.76 using RBFNN and from 380.7 to 2773.9 with RR-BLUP.

In the scenarios with h^2^=60, the MSER value improved from 4.5 to 3.4 using RBFNN and from 85.8 to 71.4 with RR-BLUP in the scenario with additive effects. In the scenario with additive dominance, the average of values changed from 5.0 to 4.0 using RBFNN and from 23.76 to 88.43 with RR-BLUP and in the epistasis scenarios the average of reliability values changed from 15.9 to 13.2 using RBFNN and from 257.9.0 to 358.3 with RR-BLUP.

## Discussion

The dimensionality reduction for the model fit is a recurring theme in several studies aimed at genomic prediction of genetic values [6,12,20,21]. However, it is worth noting that there is a difference between two approaches usually considered as dimensionality reduction ones. The first approach uses methods such as main and independent components to obtain latent variables that will be used to fit the models. With that strategy, the main goal is not to exclude markers but to use the latent variables, which are linear combinations of all available markers, to fit the model. In the second approach, the researcher has an interest in selecting the markers most related to the traits of interest and uses them in fitting the models whether they are regression ones or diversified architectures of computational intelligence [22,2,5] for their benefits both in regression models and in diversified architectures of computational intelligence. The present study considers the second approach.

In general, in terms of reliability, dimensionality reduction positively impacted all the scenarios evaluated, which represented different genetic architectures (Table 1). Better performance was not observed regarding the use of neural networks when compared with the results obtained with RR-BLUP. These results suggest that the degree of simulated epistasis, in which only dual interactions between subsequent markers are considered, was not a determining factor in differentiating the fit of regression models and neural networks. In terms of dominance, as already reported in the literature [22,23,24,25,26], that is not regarded as a problem in genomic prediction studies. Therefore, even if non-parametric models such as artificial neural networks do not need to impose strong assumptions upon the phenotype-genotype relationship presenting the potential to capture interactions between loci by the interactions between neurons of different layers [27,2], a substantial improvement in the prediction process depends on the level of epistasis present. In terms of reliability, similar results were observed in the studies carried out by [2,6], which were based on complete genome simulation with 2000 markers in a random mating population of bulls and heifers in three scenarios: additive, dominant and epistatic. In the present study, two RBFNN models were used, and in the first one there were specific weights for each SNP; while in the second one, all SNPs had the same importance. In most cases, the model with specific weights was better than that with a common weight for each SNP.

Weigel et al. [28,29] compared the use of some equally spaced markers in the genome and imputed other markers based on a reference population with all the genotyped markers using a set of markers selected according to their effect on the character of interest. The above authors concluded that when the number of selected markers is small, the predictive capacity of the model with markers selected according to the effect is higher than the use of a smaller set of markers scattered throughout the genome.

On the other hand, considering the results within the two approaches evaluated (RBFNN and RR-BLUP), dimensionality reduction also caused a reduction in the RMSE values. These results were similar to those obtained by the authors in [10], who observed that it is possible to improve prediction, both in terms of R^2^ and RMSE, predicting genetic values by means of non-parametric models when the selection includes markers that are not related to the traits of interest. When the methods were compared, a gain was observed in terms of RMSE when the fitting was performed by means of Neural Networks.

In the case of RR-BLUP, the effects of dominance and epistasis contributed to the increase of the error by increasing the difference between the expected and the observed values. In this way, when the interest is to select only a few individuals, the best 20% for example may not be the same. Similar results were observed in the study developed by the authors in [6], who used simulation of quantitative characters under different modes of gene action (additive, dominant, and epistatic) and found that RBFNN had a better ability to predict the merit of individuals in future generations in the presence of non-additive effects than by using an additive linear model, such as the Bayesian Lasso one. In the case of purely additive gene effects, RBFNN was slightly worse than Lasso. Still in the above study, the authors reported the use of the dimensionality reduction method – of the main component type – before using RBFNN and also showed that with the selection of markers the performance of the radial base network was better.

In non-parametric models, no assumption is made regarding the form of the genotype– phenotype relationship. Rather, this relationship is described by a smoothing function and driven primarily by the data. Because of that, RBFNN should be flexible with respect to type of input data and mode of gene action, such as epistasis [8,30,31,7]. This is due to the fact that artificial neural networks (ANNs) can capture non-linear relationships between predictors and responses and learn about functional forms in an adaptive manner, since they act as universal approximators of complex functions [8]. ANNs are interesting candidates for the analysis of characters affected by genetic action with epistatic effects.

Due to the importance of epistasis in studies of quantitative traits in plants [32,33,34,35,36,37], explicit (in the model) or implicit (in hidden layers) inclusion of epistatic interactions may increase the accuracy of prediction [38]. Furthermore, the frequency variation of the epistatic allele between populations may cause the gene-of-interest effect to be significant in one population but not in another, and the effect may even be inverse on the character in different environments [5], which reinforces the importance of using computational intelligence methods that easily incorporate interactions between linear effects through their hidden layers.

## Conclusion

The use of a variable selection procedure is an effective strategy to improve the prediction accuracy of computational intelligence techniques that successfully allow incorporating interactive effects, which in the present study represent biological epistatic interactions.

